# Plasmid transmission dynamics and evolution of partner quality in a natural population of *Rhizobium leguminosarum*

**DOI:** 10.1101/2024.10.17.618979

**Authors:** David Vereau Gorbitz, Chase P. Schwarz, John G. McMullen, Mario Ceron-Romero, Rebecca T. Doyle, Jennifer A. Lau, Rachel J. Whitaker, Carin K. Vanderpool, Katy D. Heath

## Abstract

Many bacterial traits important to host-microbe symbiosis are determined by genes carried on extrachromosomal replicons such as plasmids, chromids, and integrative and conjugative elements. Multiple such replicons often coexist within a single cell and, due to horizontal mobility, have patterns of variation and evolutionary histories that are distinct from each other and from the bacterial chromosome. In nitrogen-fixing *Rhizobium*, genes carried on multiple plasmids make up almost 50% of the genome, are necessary for the formation of symbiosis, and underlie bacterial traits including host plant benefits. Thus the genomics and transmission of plasmids in *Rhizobium* underlie the ecology and evolution of this important model symbiont.

Here we leverage a natural population of clover-associated *Rhizobium* in which partner quality has declined in response to long-term nitrogen fertilization. We use 62 novel, reference-quality genomes to characterize 257 replicons in the plasmidome and study their genomics and transmission patterns. We find that, of the four most frequent plasmid types, two (types II & III) have more stable size, larger core genomes, and track the chromosomal phylogeny (display more vertical transmission), while others (types I & IV – the symbiosis plasmid, or pSym) vary substantially in size, shared gene content, and have phylogenies consistent with frequent horizontal transmission. We also find differentiation in pSym subtypes driven by long-term nitrogen fertilization. Our results highlight the variation in plasmid transmission dynamics within a single symbiont and implicate plasmid horizontal transmission in the evolution of partner quality.

**IMPORTANCE:** Understanding how bacterial genes move through natural populations is critical for understanding how bacterial traits evolve. The nitrogen-fixing bacterium *Rhizobium leguminosarum* lives in symbiosis with plants and is a model for studying how gene transmission from one cell to another on mobile genetic elements called plasmids impacts the evolution of bacteria and plants. Here we characterize the genomes of a natural bacterial population, then use novel approaches to show that mechanisms of plasmid gene transmission varies across multiple plasmid types possessed by *R. leguminosarum.* We find that changes in plasmid genes are associated with the decline of symbiotic partner quality in strains isolated from environments undergoing long-term fertilization. Together, these results underscore the importance of plasmid evolution in shaping ecosystem processes like nitrogen cycling. Our study provides a framework for probing the plasmid dynamics within natural bacterial populations and how plasmid transmission affects genetic diversity and ecological interactions in bacteria.

## INTRODUCTION

Predictive models of bacterial trait evolution require a comprehensive understanding of how bacterial genes are inherited in natural populations. Bacterial traits arise and evolve via point mutations, gene duplication, homologous and non-homologous recombination, structural variants like transposons, and rare events like functionalization of non-coding regions (1). In addition, bacterial genomes can also undergo size-reduction in nutrient-limited environments or through endosymbiosis (2,3). At the core of all these processes are the vertical and horizontal transfer of genes and extrachromosomal elements (ECEs) such as plasmids, megaplasmids, and chromids (4,5) between strains. Diverse ECEs are characterized by distinct patterns of mutation, recombination, gene flow, and co-inheritance, which in turn influence how traits on these elements evolve through time (6–8). Bacterial genomes having two or more independent replicons – multipartite genomes – allow us to test how patterns of gene content, sequence similarity, size variation, and horizontal transmission rates vary across multiple extra- chromosomal elements within a single lineage of bacteria and thereby impact trait evolution.

Species with multipartite genomes are prevalent in nature and fill important ecological niches, including as opportunistic pathogens/mutualists and obligate endosymbionts. Multipartite genomes are particularly enriched in Pseudomonadota, but have been found in distant phyla including Cyanothece, Leptospira, and Deinococcus (9). The ECEs of multipartite genomes carry genes in symbionts that expand their hosts’ metabolism, as well as genes that confer resistance to antibiotics and heavy metal toxicity (10–15). ECEs can replicate independently of the chromosome, be transmitted in whole or in part between bacteria via horizontal gene transfer (HGT), and increase the size and diversity of a pangenome (the repertoire of core and variable genes found in either all, or some, members of the species, respectively) (16–19). Moreover this pangenomic variation is due in part to gene gain and loss via the action of mobile genetic elements within the pangenome, often found on ECEs (20). Due to HGT, ECEs can have evolutionary histories that are quite distinct from those of their bacterial chromosomes (21), effectively decoupling the fitness interests of ECEs and chromosomes and even driving coevolutionary dynamics among the elements within a single cell (22,23). Given the importance of ECEs in symbionts and their ability to modify their hosts’ ecology, there is a critical need for an ECE-centric approach in natural bacterial populations.

Genomic variation in bacterial pangenomes including single nucleotide polymorphisms (SNPs), indels, translocations, inversions, and duplications (24) are often concentrated on ECEs (25). Yet approaches to studying genetic variation using high-throughput shotgun sequencing tend to exclude or minimally investigate ECEs (26). For instance, common markers of phylogenetic distance such as average nucleotide identity (ANI), percentage of shared single nucleotide polymorphisms (SNPs), and shared genes are usually calculated on chromosomal genes. Reference-based assemblies fail to fully capture presence-absence diversity, and can miss major genome rearrangements, meaning that much of the genetic variation driving genome evolution can be missed or misassembled (27,28). Non-reference based, *de novo* assembly of closed, circular replicons is made difficult by repetitive elements in bacterial genomes and particularly ECEs, resulting in genome fragmentation (29) and hindering the assembly and analysis of ECEs and pan-genomes (30,31). Harnessing long-read technology to assemble populations of closed, reference-quality genomes enables pan-genome analysis of genetic diversity in natural populations of ECEs – the diversity on which selection acts.

Nitrogen-fixing rhizobia, bacteria that fix nitrogen in symbiosis with leguminous plants (32), are good models for the evolutionary ecology of ECEs since they contain multiple and diverse plasmids in their multi-partite genomes and because the genes required for associating intracellularly with hosts are found on plasmids or Integrative and Conjugative Elements (ICEs) (7,33,34). The *Rhizobium leguminosarum* species complex, formerly all one polyphyletic species, was recently divided into at least 5 genospecies (35); most recently, certain genospecies (*gsA, gsB, gsD*) have been classified as new species (*Rhizobium brockwellii*, *johnstonii,* and *beringeri,* respectively) (36). *Rhizobium* isolates are well-known to carry many plasmids, which can coexist in the same cells due to variation at the *repABC* operon (35,37); *repA* in particular determines incompatibility group, and the same *repA* “Rh group” has been found across multiple genospecies (35). These diverse *Rhizobium* plasmids tend to contain non-essential genes and are mobile, based on studies showing that the phylogenetic histories of plasmid genes are distinct from those of the chromosome (38). Fully-resolved genomes for many natural isolates are required to address variation in phylogenetic history within and among plasmids and thus understand plasmid transmission patterns on timescales relevant to bacterial trait evolution.

Clover-associated *Rhizobium* are a particularly powerful tool for understanding the contribution of plasmid diversity and plasmid transmission to bacterial evolution in nature. The canonical symbiosis genes that enable *Rhizobium* to interact with clover, vetch, and other hosts are not only found on plasmids, but have been found on *different* plasmids among natural isolates, indicating HGT of this critical gene region across replicons (32,35,39,40). In our previous work using a population of *Rhizobium* from the Long Term Ecological Research (LTER) site at Kellogg Biological Station (KBS) in Michigan, we found that strains from old field communities exposed to long-term nitrogen (N) since 1998 are less-beneficial on average for clover (*Trifolium spp*.) host plants, compared to control (41). Moreover we used reference-based SNP analysis to associate this decline in partner quality, *i.e.,* the benefits that symbionts provide to their host, with differentiation at the canonical symbiosis gene region (42). Beyond contributing to fundamental knowledge on plasmids, a better understanding of plasmid inheritance in these multi-partite genomes is critical for understanding symbiosis gene transmission and thus the role of HGT in symbiosis trait evolution.

In this study, we generate high-quality reference genomes to study ECE dynamics in a natural bacterial population, leveraging 62 previously-studied strains of *Rhizobium* (41–43). We use this population to understand how sequence diversity, gene function, and size differ among plasmids within a pan-genome (the plasmidome). We then use phylogenomic approaches to infer how the propensity for horizontal and vertical inheritance differs across plasmids in the pan- genome. Finally we study the pSym to examine the role of plasmid HGT in partner quality decline.

## RESULTS

### Delineating the plasmidome

To assess the ECE diversity in natural populations of *R. leguminosarum*, we used long-read technology to sequence and assemble complete genomes *de novo* for all 62 nodule isolates from Weese et al. (41). Each strain’s assembled genome contained 1-6 extrachromosomal elements. In total, we identified 257 extrachromosomal elements in our bacterial population. Of those, 256 had at least one *repABC* operon, which has been used as a basis for plasmid categorization (35,38,44,45), and 17 replicons had two distinct *repABC* operons. One 1.4 Mbp replicon in strain 773_N did not contain identifiable plasmid replication genes.

We first reconstructed a phylogenetic tree from a core chromosome alignment (Fig. S1), which showed that 56 of our 62 strains formed a single clade identified as *R. leguminosarum* sensu stricto, or *genospecies E* (*gsE*) (35). Four of the 62 strains (061_N, 173_C, 209_N, 231_N) were identified as *gsB*/*Rhizobium johnstonii* (36), and were characterized within our population by low diversity in the chromosomes shared between these four strains. The remaining two strains, 717_N and 773_N, fell outside both *gsE* and *gsB* clades; strain 717_N is sister to the *gsB* clade, and strain 773_N is the most basal strain. To further investigate the identity of strain 773_N, we queried NCBI non-redundant database with its 16S rRNA and *dnaA* sequences and found that the closest sequence identity match with a complete genome for both sequences was *Rhizobium sp.* WYJ-E13 (accession ID: PRJNA738292), 99.12% and 92.86% respectively.

To categorize all plasmids into discrete groups, we used a k-mer approach that considers core and non-core regions of all plasmids. We calculated the k-mer signatures for all 257 replicons and generated a weighted undirected network component graph based on the Jaccard Index for all-vs-all combinations (Fig 1). This approach benefits from using all genetic information in both the core and non-core regions of the plasmid, as opposed to using only core gene-coding regions present in all plasmids. This more inclusive approach is especially important since there is little overlap in genes across all replicons. Of 257 plasmids, most (226) fell into one of four plasmid clusters (hereafter type I, II, III, and IV; Fig. 1), while the few remaining plasmids fell into an additional four types (hereafter type V, VI, VII, and VIII).

**Figure 1:**
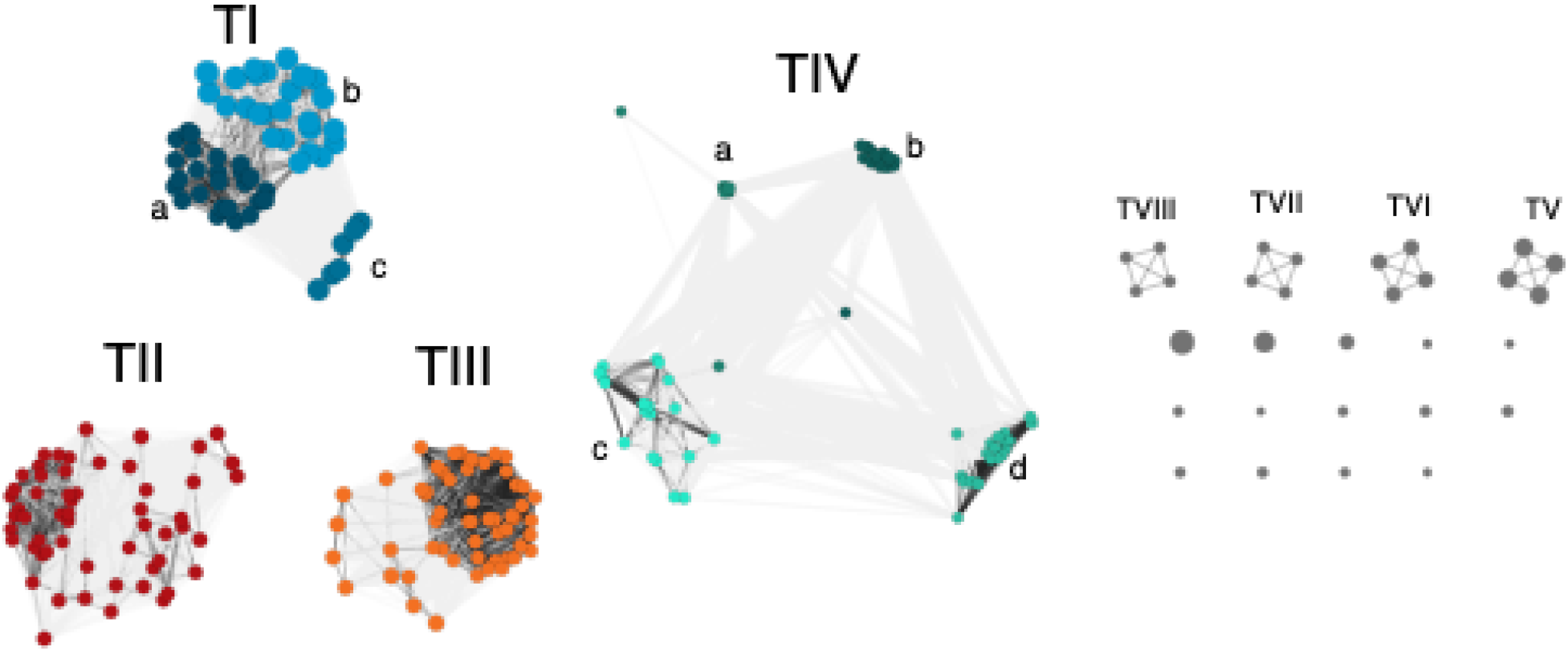
Weighted, undirected network of all 257 plasmids from a population of 62 clover- associated natural isolates of *Rhizobium.* Plasmids (nodes) were grouped by Jaccard-Index similarity > 0.1 into types. Darker edges indicate more pairwise similarity between nodes. Nodes were colored blue (type I), red (type II), orange (Type III), and green (Type IV) and shaded by subgroup for type I and IV plasmids. Plasmid types with few representatives (V-VIII and singletons) were left unshaded.

Plasmids were numbered based on sample number, and then size within the population. Both Type I and Type IV plasmids clustered into 3 and 4 smaller sub-groups, respectively (labeled a, b, c, d; Fig. 1)

The distribution of these plasmid types in our study correspond to the bifurcations in the chromosome phylogeny (Fig. 2), with different genospecies possessing distinct plasmidomes.

**Figure 2:**
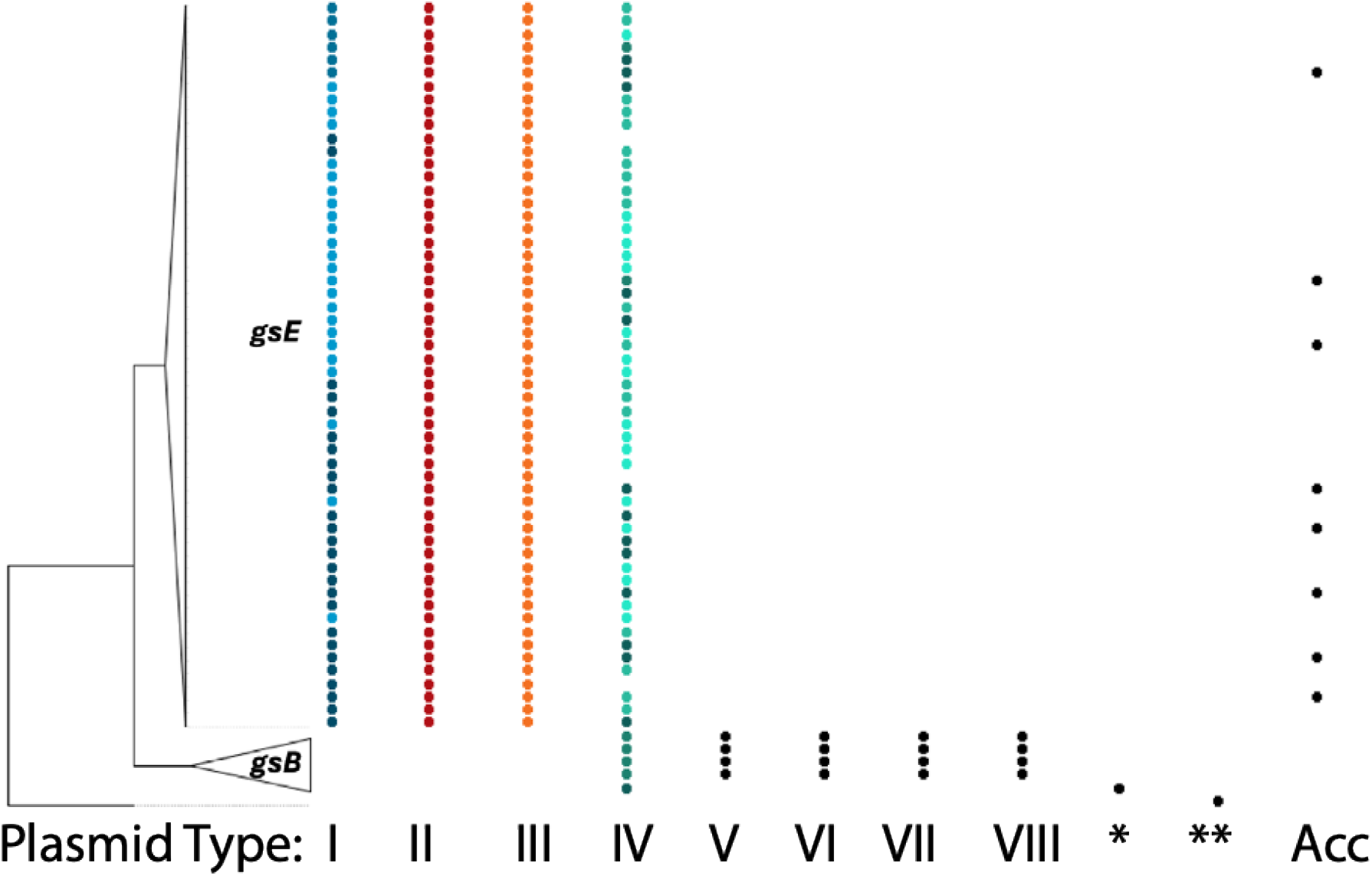
Distribution of plasmid types across the concatenated core chromosomal tree from a natural population of 62 clover-associated strains of *Rhizobium*. Tree tips were collapsed to highlight differences in plasmid composition between genospecies (*gsE* and *gsB*). Subclades within types are represented by shape for Type I: circles (Type I-a), squares (Type I-b), and triangles (Type I-c) and Type IV: circles (Type IV-a), squares (Type IV-b), triangles (Type IV- c), and diamonds (type IV-d). Six replicons from strain 717_N (*) and a single 1.4 Mbp replicon from strain 773_N (**) are each represented by a single dot.

Type IV plasmids were found in 58/62 strains across both genospecies (*i.e., gsB* and *gsE*); type I, II, and III plasmids were present in all 56 *gsE* strains (Fig. 2), while the plasmid type groups V, VI, VII, and VIII were present in all four *gsB* strains, indicating that plasmid types delineated by k-mer analysis track with chromosomal genospecies. Like their associated *gsB* chromosomes, Type V-VIII plasmids contain little genomic diversity (table 1).

**Table 1:** Summary of *Rhizobium* plasmids featured in this study. For plasmids with more than one representative, we report the range of plasmid size, core length, number of core genes, nucleotide diversity (π), and average nucleotide identity (ANI). Rh type is based on *repABC* sequence as in previous studies (35).

| Plasmid Type | N | present in | Size (Mbp) | Core length (Mbp) | Core genes | $\pi$ | ANI | Rh Type |
| --- | --- | --- | --- | --- | --- | --- | --- | --- |
| I | 56 | gsE | 0.92-1.43 | 0.62 | 510 | 0.015 | 97.94 | 1 |
| II | 56 | gsE | 0.51-0.63 | 0.39 | 335 | 0.02 | 97.32 | 2 |
| III | 56 | gsE | 0.56-0.73 | 0.50 | 421 | 0.017 | 97.89 | 4 |
| IV | 58 | gsE, gsB, Rht_717_N | 0.26-0.43 | 0.05 | 49 | 0.021 | 95.25 | 4, 6, 7, 8 |
| V | 4 | gsB | 0.93 | 0.93 | 587 | 0 | 100 | 1 |
| VI | 4 | gsB | 0.67 | 0.67 | 419 | 0 | 100 | 2 |
| VII | 4 | gsB | 0.39 | 0.39 | 219 | 0 | 100 | 5 |
| VIII | 4 | gsB | 0.35 | 0.35 | 246 | 0 | 100 | 3 |
| IX | 1 | Rht_717_N | 1.10 | - | - | - | - | 1 |
| X | 1 | Rht_717_N | 0.61 | - | - | - | - | 2 |
| XI | 1 | Rht_717_N | 0.31 | - | - | - | - | 3 |
| XII | 1 | Rht_717_N | 0.29 | - | - | - | - | 9 |
| XIII | 1 | Rht_717_N | 0.21 | - | - | - | - | 4 |

We next examined the presence of the canonical symbiosis genes (*i.e. nif, fix,* and *nod*) in our annotated genomes, and found they were limited to type IV plasmids (pSym hereafter).

Therefore, the presence of type IV plasmids appears to be required for symbiosis with clovers (*Trifolium* spp.) in this population of *Rhizobium*. Interestingly, we found that four of the 62 strains lacked the pSym, further corroborating that this non-essential element could be lost in strains present within a naturally occurring population (42). Given that all 62 strains were isolated from nodules, it is not known whether this loss occurred in culture or whether these isolates co-infected nodules with symbiotically-capable strains (46,47).

Of the remaining smaller clusters of plasmids, fourteen unique plasmids did not group with any other plasmid within the population (Fig. 1). Of these singleton replicons, six were found in strains outside the main *gsE/gsB* clades; five of the singleton replicons belonged to strain 717_N (type IX, X, XI, XII, and XIII), and one belonged to strain 773_N (Fig. 2). The other eight singleton plasmids (hereafter, accessory plasmids) were in strains scattered across the *gsE* phylogeny (Fig. 2).

### Genomic content of the plasmidome

To visualize the difference in gene composition between plasmids, we annotated the genomes using NCBI’s PGAP (48), assigned genes to clusters of orthologous genes (COG) groups (49), and performed a Principal Coordinate Analysis (PCoA) with the Jaccard distance calculated from the presence-absence of orthologous gene clusters of the plasmidome. We found that, using PCoA based on orthologous gene content, all 257 plasmids formed clear clusters that correlated with plasmid types from k-mer clustering (Fig. 3), further indicating that these plasmid types differ in gene content and have distinct functional roles. In the PCoA, the pSyms cluster together in the center between plasmids I-III, and alongside the accessory plasmids, suggesting some shared gene content among these elements. In particular, many 209) genes were shared between at least one accessory plasmid and one Type IV plasmid.

**Figure 3:**
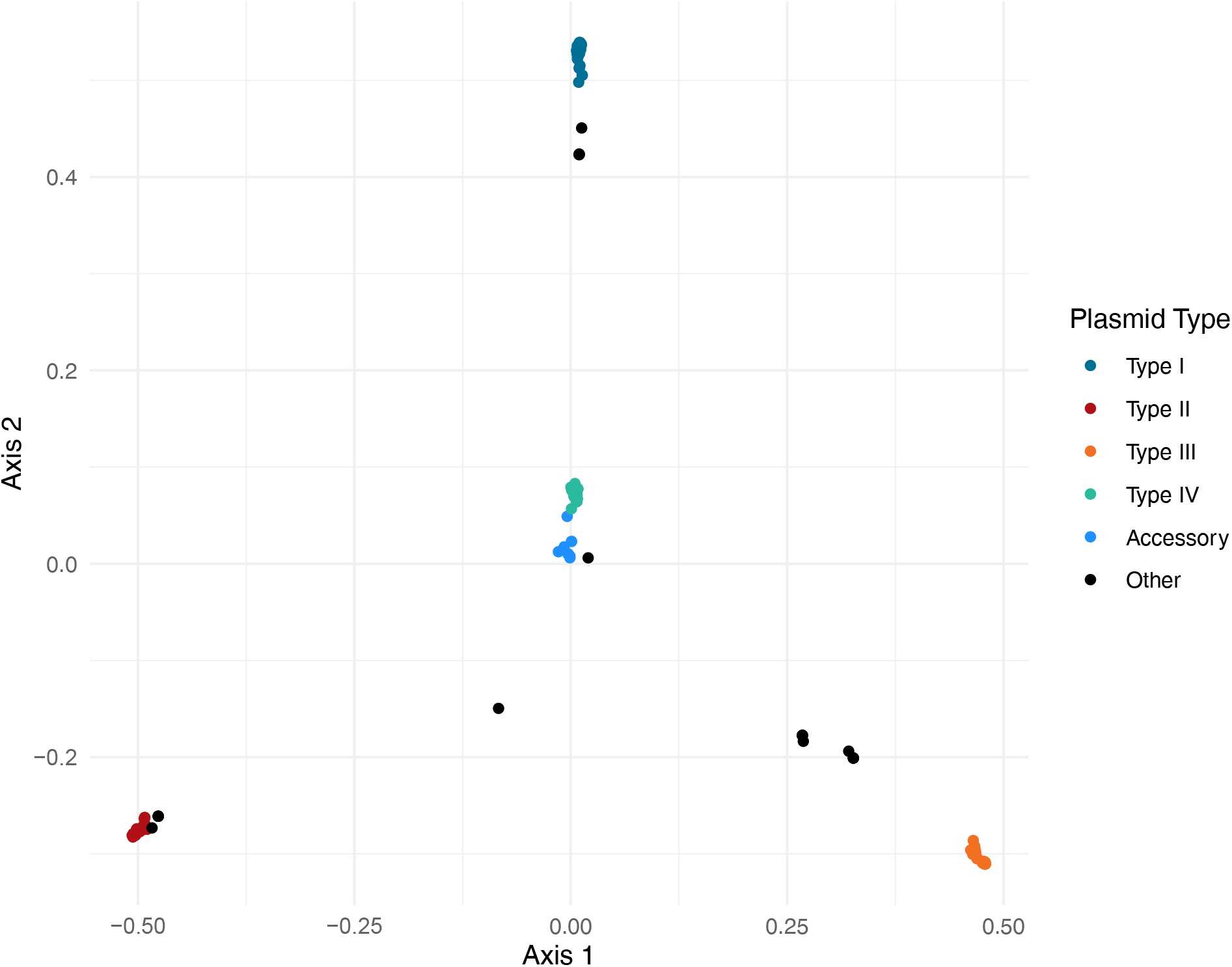
Principal Coordinate Analysis (PCoA) based on the presence-absence matrix of orthologous gene clusters in the *Rhizobium* plasmidome of 62 natural isolates. Each point represents a single plasmid, with points closer together indicating more shared gene content.

Although the *gsB* plasmid types V-VIII formed k-mer clusters distinct from *gsE* plasmids (Fig. 1), these four plasmids were similar to (had low Jaccard distances with) *gsE* plasmid types I (V), II (VI), and III (VII and VIII) (Fig S2A), suggesting that these sets of elements serve somewhat analogous functions in *gsB* and *gsE*. Indeed pairwise comparison of type I with V and type II with VI indicated 56% and 74% core gene content overlap, respectively (Fig. S2B).

Interestingly, while pairwise distances between *gsE* plasmid type III and either *gsB* type VII or VIII were low (Fig. S2A), pairwise distance between *gsB* types VII and VIII was very high (Fig. S2B); these patterns suggest that the core gene content of type III in *gsE* is comprised of a combination of the cores of Types VII and VIII in *gsB*. This is supported by an alignment of types VII and VIII to the type III plasmid, showing large syntenic aligned blocks consistent with shared plasmid history (Fig. S2C). Interestingly, while plasmid types I and V were both in Rh group 1, and II and VI were both in Rh group 2, plasmid types III, VII, and VIII belonged to three different Rh groups (4,5, and 3, respectively) (Table 1) – indicating that plasmid gene content often, but not always, tracks *repABC*-based Rh group.

Overall, COG analysis suggested that high-level distribution of functional gene content was similar across all plasmids (Fig. S3), despite their unique patterns of gene presence-absence variation (PAV). Across all plasmid types in the population, four COG categories were completely absent in the functional prediction (A, B, W, Y). The absence of these functions was not surprising as they relate to RNA processing (A), chromatin (B), extracellular (W), and nuclear (Y) structures, which are more commonly associated with eukaryotic or chromosomal processes. The one outlier was the pSym, which features more genes for intracellular trafficking and secretion (U), and recombination and replication (L) (Fig. S3).

### Variable size distributions in gsE Rhizobium plasmids

Because we have the largest sample of plasmids from *gsE* strains (I-III, and pSym), hereafter we primarily focus on the plasmidome of the *gsE* subgroup in order to examine within-type variation and transmission. Plasmid types varied in size and in the shape of their size distribution (Fig. 4). Type I plasmids were the largest in our population, had the largest size variation, and had a bimodal distribution centered at either ∼0.9 Mbp or ∼1.2 Mbp. Type II and type III plasmids were similar in size, with intermediate lengths ranging from ∼0.5-0.6 Mbp. Of the *gsE* plasmidome (types I-IV), pSyms were the smallest (∼0.2-0.4 Mbp). Notably, this symbiosis gene location is distinct from the reference genome WSM1325 (*gsA*), in which the symbiosis genes are found on the largest plasmid (32).

**Figure 4:**
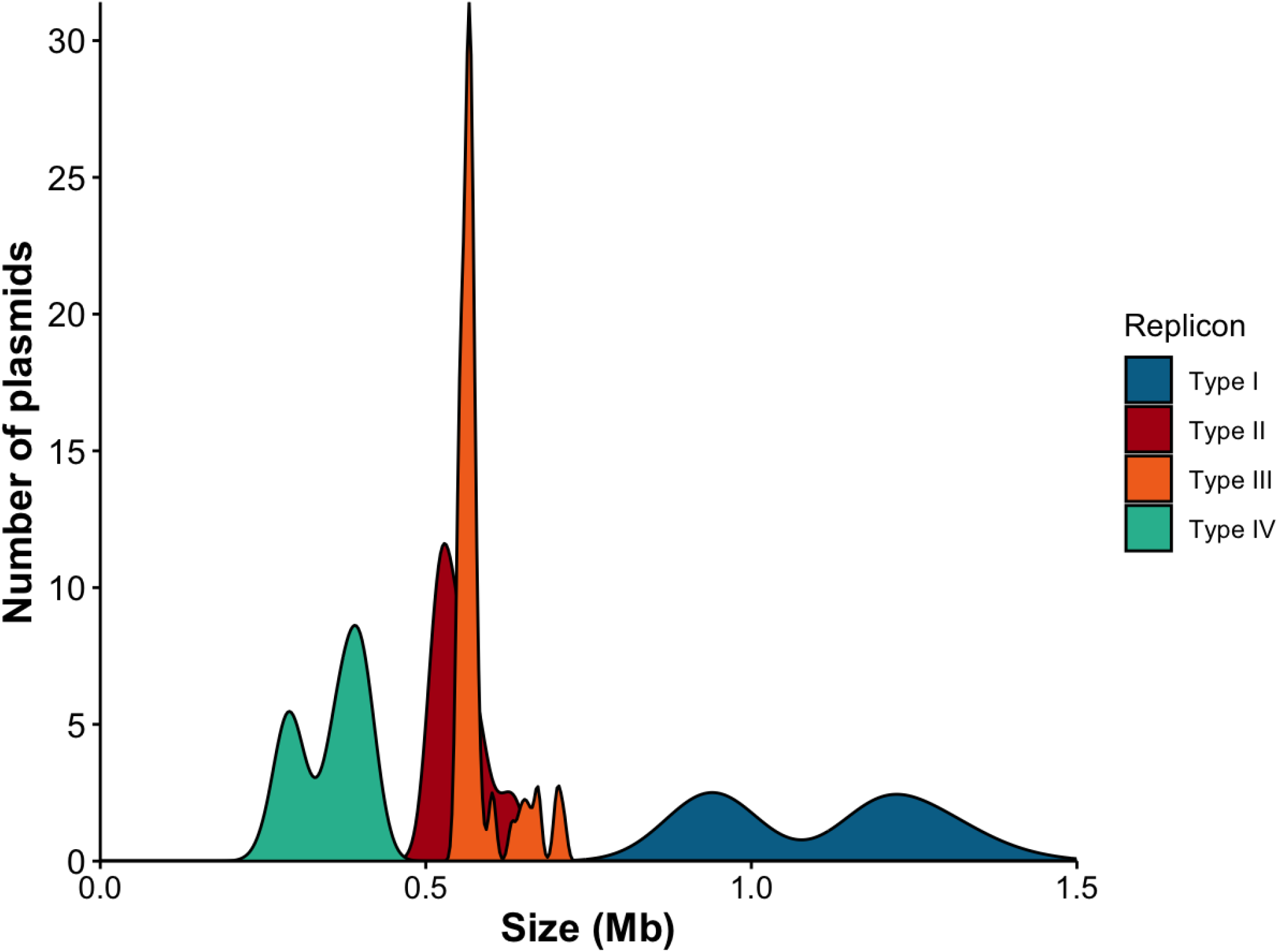
Distribution of plasmid size for each of the four major plasmid types (I-IV) commonly found in *Rhizobium* genospecies (*gsE*). Distributions of types I-III include plasmids from all 56 *gsE* strains, whereas the type IV distribution includes 58 type IV plasmids from *gsE* and *gsB* strains, as well as 717_N and 773_N.

Comparing the size variation within the type I plasmid with its core length, content, and phylogeny (Fig. 4, Table 1) allows us to reconstruct the history of how dramatic changes in plasmid size have evolved in this element. The three type I k-mer subgroups (I-a, I-b, I-c; Fig. 1) map onto the three distinct groups in the type I core phylogeny (Fig. S4a). However although subgroup I-b and I-c have similar size distributions, they contain distinct large insertions (Fig. S4b). All small type I plasmids (subgroup I-a) form a clade that diverged from I-b after a single loss of the ∼0.3 Mb insertion (Fig. S4b, c). Thus independent gain and loss of large insertions is responsible for the bimodal size distribution; in addition, these large insertions occur in similar locations along the plasmid (Fig. S4c), suggesting that type I plasmids might have specific hotspots where insertions are more likely.

### Mode of inheritance varies in the Rhizobium plasmidome

To study plasmid transmission modes, we used a phylogenetic approach with random resampling of orthologous genes to quantify patterns of gene tree heterogeneity within each plasmid type. We then calculated the Generalized Robinson-Foulds (GRF) distance between two trees as a measure (0–100) of phylogenetic congruence between two trees. Mean GRF distance between gene trees of the chromosome was 72.46, with a standard deviation of 5.40 (Fig. 5). The type I and type II plasmids had similar distributions to each other and to the chromosome (mean = 72.84 ± sd 5.08 and 68.00 ± sd 6.31, respectively) – suggesting similar levels of within-element vertical versus horizontal transmission on these three elements. By contrast, the distribution of type III plasmid GRF distances (mean = 56.41 ± sd 5.09) indicates that gene trees within this element were more similar – suggesting more internal consistency and thus less horizontal transmission of genes compared even to the chromosome. Finally, the pSym GRF distribution was wider and centered at mean = 86.79 ± sd 9.46, indicating higher divergence of gene trees among the loci on this element, compared to the other four – consistent with abundant HGT of genes on the pSym.

**Figure 5:**
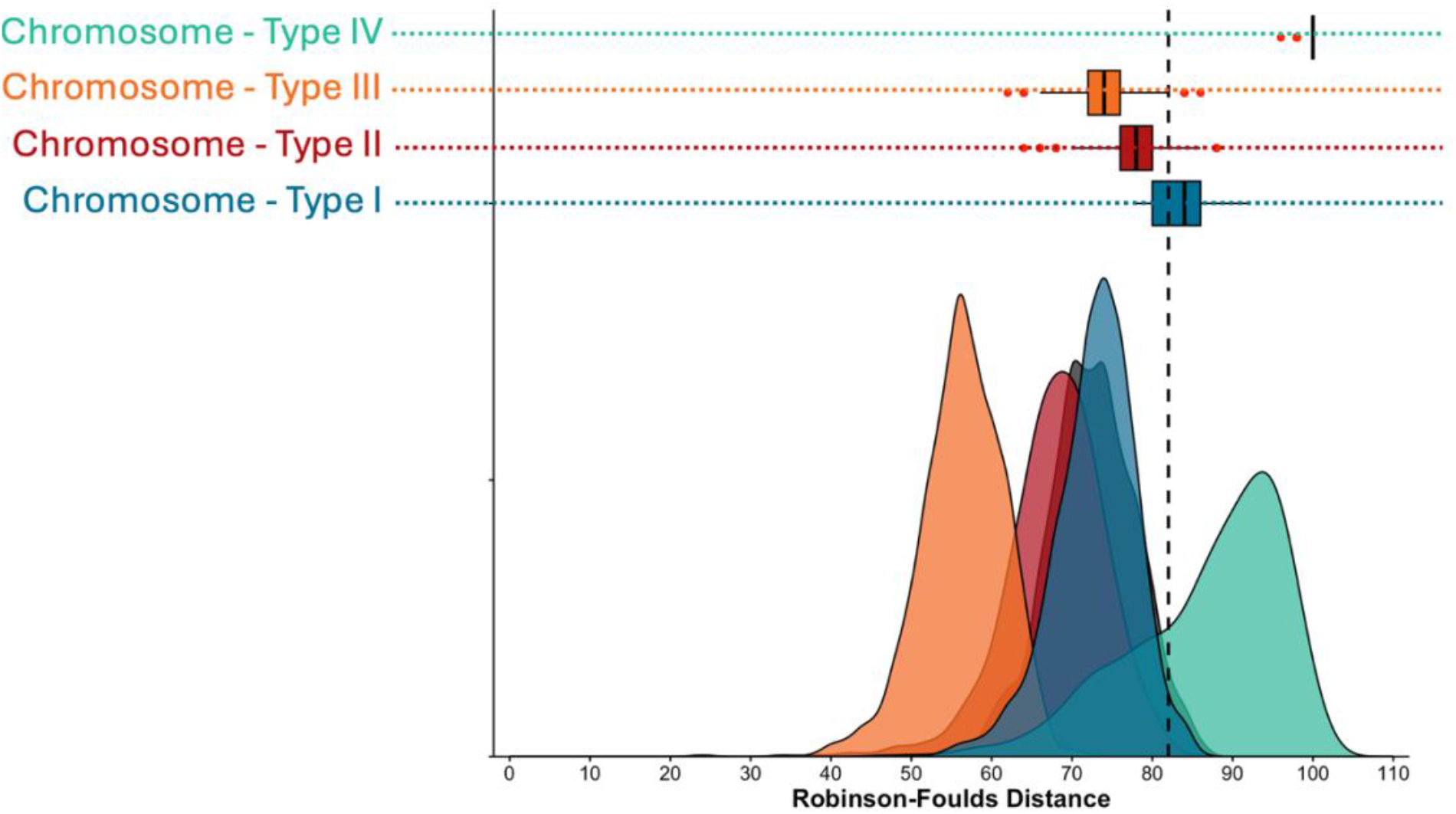
GRF distance distributions of within-element and across-element gene trees. Values closer to 0 represent higher levels of across-tree congruence, while values closer to 100 represent higher levels of incongruence among trees. Within-element distances for the chromosome (gray), Type I plasmid (blue), Type II plasmid (red), Type III plasmid (orange), and Type IV plasmid (green) are shown as geometric smoothed distributions along the bottom. GRF distance distributions between each plasmid and the chromosome are represented as horizontal box and whisker plots. The vertical dotted gray line represents the 95^th^ percentile of the within- chromosome distribution.

Next, to assess the degree to which plasmids are vertically inherited (together with the chromosome), versus horizontally (separately from the chromosome), we compared the distributions of GRF distances when gene trees from each plasmid were compared to those from the chromosome. The distributions of GRF values for plasmid types II and III fell inside the 95^th^ percentile of the chromosome GRF distance distribution (Fig. 5), indicating that the likelihoods of co-inheritance of chromosomal genes with genes on these plasmids were not different from the co-inheritance of chromosomal genes with each other – supporting abundant vertical transmission alongside the chromosome in this population. By contrast, the chromosome-type I gene distribution (mean = 83.24 ± sd 2.97) fell above the 95^th^ percentile of the chromosomal gene distribution (Fig. 5), indicating that the genes on these two elements tend to have distinct evolutionary histories, suggesting HGT. Finally, GRF distances between genes on the chromosome and the pSym were particularly high (mean = 99.56 ± sd = 0.83, with 881/1000 resamplings resulting in a maximum GRF = 100; Fig. 5), also suggesting high levels of horizontal transmission of pSym genes relative to the chromosome – likely due to the HGT of entire pSym plasmids across chromosomal lineages. Due to the small core, robust clade patterns, and high levels of within-subclade core genes of the pSym (see below Fig. 6A,6B), we separately ran the resampling analysis of gene tree heterogeneity by pSym subclade to test whether these patterns of HGT result from across-clade differences rather than individual plasmids moving independently of the chromosome. Despite high sequence similarity within Type IV-a, b, c, these subclades show elevated GRF distances compared to the chromosome (Fig. S5A), suggesting that the entire pSym is moving horizontally.

**Figure 6:**
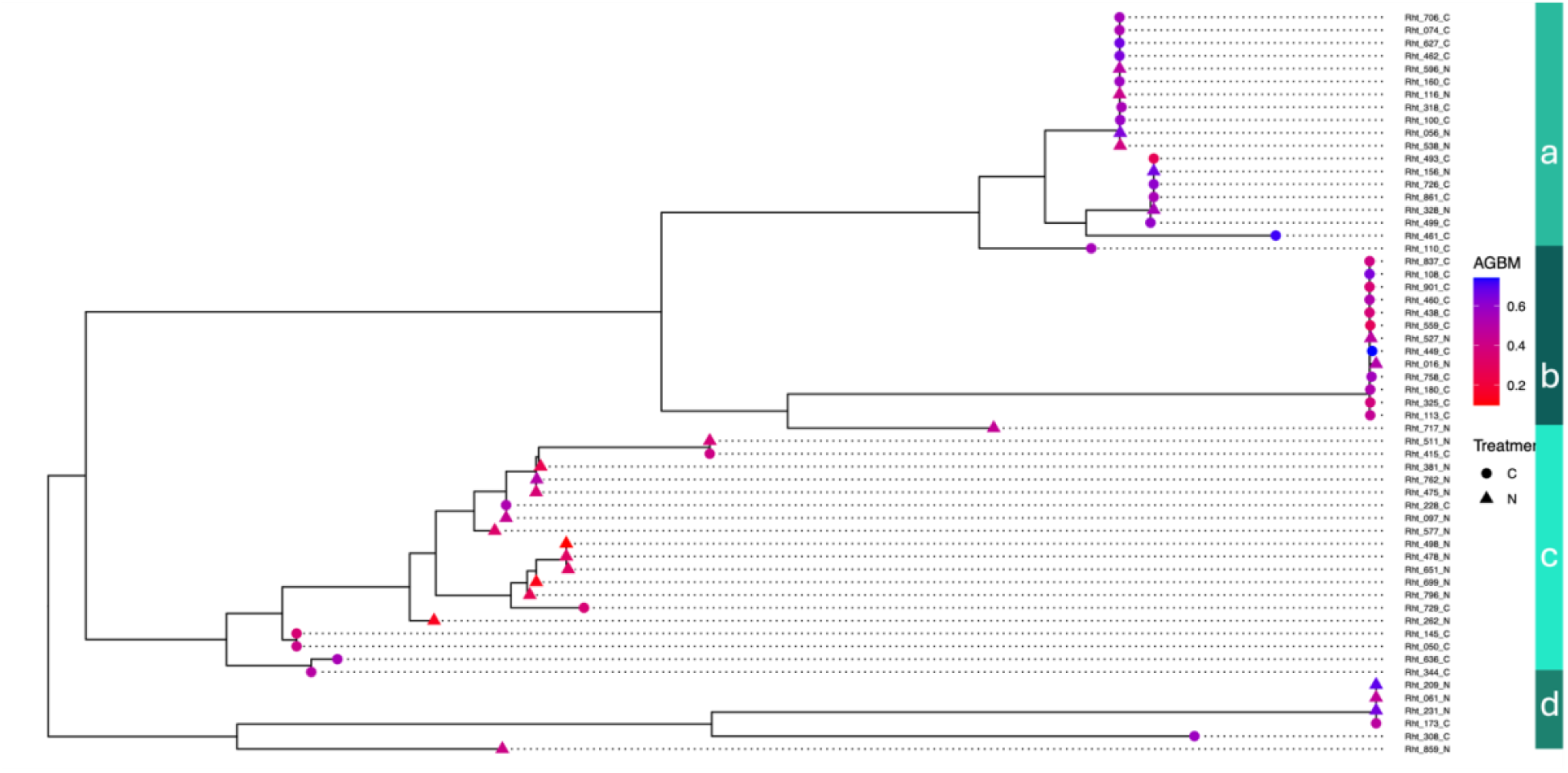
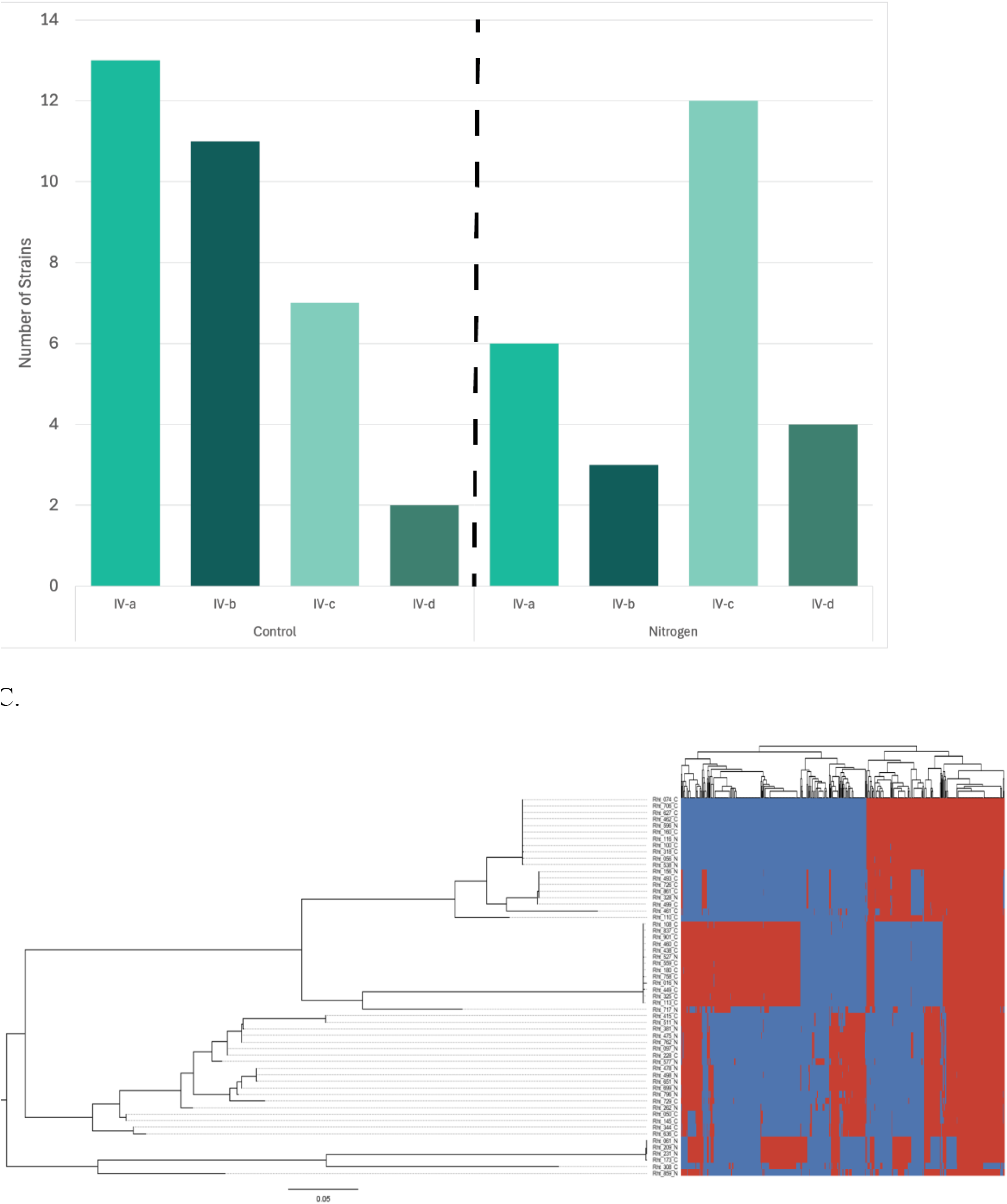
(A) Phylogenetic tree of the type IV plasmid (pSym) based on concatenated core gene alignment, with nodes labelled with aboveground biomass (AGBM as color gradient) and plot treatment of origin (C for control or N for N-fertilized). Type IV subclades (a-d) as in Figure 1 are indicated in shades of green on the right. (B) Number of strains from each Type IV sub-clade isolated from either control (left) or N-fertilized (right) plots from the KBS LTER. Any nodes with < 85% bootstrap support were collapsed.

### pSym (Type IV) plasmid movement drives differences in partner quality

Given the evidence for abundant HGT of the pSym plasmid, and previous population genetic analyses focusing on symbiosis-related loci (42), we next studied patterns of differentiation in the pSym in order to relate pSym variation to symbiotic partner quality. First, we inferred a phylogenetic tree with concatenated core sequences from the type IV plasmid and found four clades (Fig. 6A); these clades mirror the four major k-mer clusters of type IV plasmids (Fig. 1).

The pSym is characterized by a particularly small core (52.1 kb, or 49 genes) relative to other types (Table 1). Although a gene presence-absence plot shows modularity and unique gene content unique in subclades IVa-c (Fig. S5B,C), groups of shared genes are often found in unrelated clades (Fig. S5C), and the core gene set of any two pSym subgroups (regardless of phylogenetic distance) is noticeably larger than the universal pSym core of 49 genes – indicating abundant gene-sharing across groups on the pSym phylogeny (Fig. S5C, D). For example, despite being more phylogenetically distinct and most often found in the *gsB* minor clade, nearly all IV-d genes are shared with other type IV sub-clades (Fig. S5C).

We found clear cases of pSym HGT across distinct chromosomal lineages in our population. First, although strain 717_N is more closely related to *gsB* than *gsE* at the chromosome (Fig. S1) and shares no other plasmids with either group (Fig. 2), the 717_N pSym falls alongside a group of high-quality *gsE* strains in the pSym tree (clade IV-b, Fig. 6A).

Similarly, although the *gsB* and *gsE* chromosomal clades generally have distinct plasmidomes including the pSyms (Fig. 2, Fig 6A), two *gsE* strains (308_C and 859_N, Fig. S1) were found to carry *gsB*-like pSyms (clade IV-d; Fig. 6A) indicating cross-genospecies HGT.

Finally we found striking correspondence between the core pSym phylogeny and the benefits of symbiosis for plant hosts, as measured in a previous common garden experiment (41). Within the major pSym groups, clades IV-a and IV-b contain mostly higher-quality strains that originated from unfertilized control plots (Fig. 6A), whereas clade IV-c contains an abundance of lower-quality strains isolated from N-fertilized plots and features longer branch lengths due to SNPs in core genes (Fig. 6A). A Fisher’s exact test supported a significant difference in frequency of Type IV sub-clades between control and N treatments (p = 0.042; Fig. 6B).

## DISCUSSION

Plasmid inheritance is a key process underlying bacterial trait evolution in natural and managed ecosystems. Studies of plasmid variation and transmission within local-scale, recombining populations are needed to quantify patterns of gene co-inheritance as well as their influence on bacterial phenotypes. Here we delineate the major types, size, gene content variation, and transmission patterns of coexisting plasmids from a single population of clover-associated *Rhizobium*. We find that *Rhizobium* plasmids vary considerably in their size structures and modes of transmission; some plasmids (type II, type III) appear to be primarily vertically transmitted, while others (type I, pSym) are more likely to be horizontally transmitted.

Concomitant with these findings from within our best-sampled genospecies (*gsE*), we find that most of the plasmidome is delimited by chromosomal lineages. Nevertheless, the extent of this limitation varies across plasmids; for example, we find clear examples of cross-genospecies HGT of the pSym. Finally, our analysis of pSym subclades indicates a role for pSym HGT in the decline of clover-associated partner quality in N-fertilized environments. Below we discuss the important lessons learned from each of the major element in our *Rhizobium* plasmidome, then finish with a holistic discussion of the potential importance of transmission variation to the evolution of bacterial traits.

### Type I plasmids

The largest plasmid in *Rhizobium* frequently carries the symbiosis genes (42,43). Indeed in previous work (42) using referenced-based assembly to WSM1325, we assumed that the Type I plasmid was the symbiosis plasmid (see discussion of the pSym below). This highlights the value of using non-reference-based genome assembly facilitated by long read sequencing, as well as naïve k-mer based clustering, for studying plasmid variation, plasmid transmission, and the evolution of plasmid-borne traits. Plasmid size variation is usually caused by differing patterns of presence and absence of genes, which is caused by homologous recombination or horizontal gene transfer (50,51). Studies of plasmid evolution usually limit analyses to a handful of genes involved in plasmid replication, maintenance and transfer (52,53). The ability to interrogate fully closed plasmid genomes allowed the separation of core and variable content within this single plasmid type, revealing the gain and loss of large insertions that dramatically alter plasmid size and track the core phylogeny. The internal consistency of the type I plasmid (based on the comparison of core and variable content, and similar *within*-plasmid gene tree distances to those of the chromosome) at first appears at odds with evidence for HGT *between* type I plasmids and the chromosome. These results might hint at distinct mechanisms governing within-element stability versus whole-element transfer in this plasmid (*e.g.,* homologous recombination versus conjugation), though more functional studies are required to make any generalizations.

### Type II and III plasmids

We found that the type II and type III plasmids are more frequently vertically transmitted alongside the chromosome, compared to the other plasmids, and accordingly, have a large fraction of core genes. In contrast with type I and IV plasmids, the gene PAV in type II and III plasmids is comprised of singleton genes being present at low frequencies in the plasmids as opposed to large gene-clusters being present in multiple closely related plasmids. This suggests that the mechanisms by which type II & III, and type I & IV are acquiring and losing genes are different. In fact, the type III plasmid shows particularly low *within*-plasmid GRF distances, suggesting particularly low rates of recombination, though the potential mechanisms remain unclear.

Our observation that the *gsE* type III plasmid contains the gene content of *two* other plasmids from the *gsB* group (type VII and VIII) might suggest a key evolutionary event in the history of these *Rhizobium* plasmids – either a subdivision of one (fission) or a merging of two (fusion) – and denotes a key change in how *gsE* and *gsB* strains subdivide their respective genomes. Importantly, however, the type III does not share its Rh incompatibility group (35) with either of these plasmids, making historical reconstruction difficult. Chromosomal fusion is known in closely-related *Agrobacterium tumefaciens* (*54*). The “schism hypothesis” of chromosome fission has been developed as an explanation for the evolution of multi-partite genomes (51); such processes might generate plasmid diversity as well. Fusion and fission of eukaryotic chromosomes is well-known to drive reproductive isolation and thus speciation in plants and animals (54). Understanding the impact of these processes in the diverse plasmids of *Rhizobium* and other species is paramount to further understanding bacterial genome evolution and the diversification of bacterial species.

### Type IV plasmids

The pSym is usually defined as the replicon that harbors genes necessary for symbiosis with a host plant (23). In our population, the pSym is the type IV plasmid, the smallest of the main plasmids and the only one present in both *gsE* and *gsB* chromosomal genospecies. It is also the most variable, having a very small pool of core genes and deeply diverged lineages.

Nevertheless, we treat the pSyms as one “type” of plasmid for multiple reasons. First is their clustering based on the k-mer approach we used to categorize plasmids. Though previous work has used *repABC* Rh types to categorize *Rhizobium* plasmids (35), we found that 17 pSyms in our population contain two distinct full *repABC* operons, suggesting the need for an additional approach that reflects gene content similarity. Because plasmids with the same *repABC* groups generally cannot be maintained in the same cell (55), it is unclear what the evolutionary advantage of having two distinct copies of the operon in a single plasmid might be. Second, while some pSym lineages possessed subclade-specific gene clusters, there was abundant shared gene content in pairwise comparisons of the pSym subclades regardless of relatedness at the core – suggesting mobility of pSym genes across the pSym phylogeny. Finally, examining all pSyms together based on gene content and function allowed us to relate pSym subclade to symbiotic partner quality and detect shifts in subclade frequency between N-fertilized and control plots (see below).

The symbiosis plasmids in the *R. leguminosarum* species complex are well-known to exhibit high levels of variable gene content and horizontal mobility (23,56,57). Nevertheless we found that the pSym subclades tended to be associated with different chromosomal genospecies (*gsE* and *gsB*), and the few notable exceptions allow us to pinpoint clear cases of pSym HGT across the genospecies boundary. Previous approaches designed to detect introgression events across genospecies support shared alleles at symbiosis genes *nifB*, *nodC*, and *fixT* across *gsE* and *gsB* (35); our results suggest that the symbiosis plasmids move across these genospecies boundaries as well. Although a much more thorough functional analysis would be required to pinpoint the underlying drivers of transmission in our plasmids, extra recombination genes (L) in the pSym might explain higher levels of recombination and PAV in this plasmid. Nevertheless it is interesting to consider the genetic drivers and selective forces that might reinforce, versus break up, this type of structure in chromosome-plasmid relationships through time and across environmental conditions (*e.g.,* the presence or absence of hosts, changing abiotic conditions (51,58)).

Previously, we had reported differentiation of the symbiosis gene region between high-quality partners in the control plots and low-quality partners in the N-fertilized plots in this *Rhizobium* population, relative to the rest of the genome (42). Here, we add evidence that it is the entire pSym, and not just the symbiosis gene region, that is differentiated. Together with evidence for HGT of the pSym – both between chromosomal genospecies and within a single, well-sampled genospecies (*gsE*) – we infer a shift in pSym subclade frequencies with a change in the environment, rather than a gene-specific sweep at the symbiosis gene region. The shorter branch lengths in higher partner quality pSym subclades might indicate purifying selection in control plots, or relaxed selection in N-fertilized plots. Our new, plasmid-centric interpretation of the genetic underpinnings of partner quality decline stems from both data type and analytical methods; our long read-enabled full genome assemblies reveal diversity in symbiosis gene location, duplicate pSym *repABC* types, and pSym gene content that was not previously visible. What’s more, these fully-resolved plasmidome sequences, combined with novel phylogenomic analyses, allow us to quantify patterns of gene tree heterogeneity not only at the pSym, but across all plasmid types and thus make inferences about the variation in transmission modes among elements within a single genome.

### Transmission in the Rhizobium plasmidome

The degree to which plasmid vertical versus horizontal transmission modes are determined by chromosomal mechanisms, plasmid-specific mechanisms, and/or the interaction is still being worked out (45,59). It has long been recognized that plasmid transmission mode can evolve as the costs and benefits of conjugation-growth tradeoffs shift (60,61), though the selective drivers in nature are not well-known. Most plasmid transmission studies take place in the lab under strongly selective conditions in one or a few laboratory or clinical strains (62,63), whereas population genomic studies on natural diversity have historically not focused on ECEs given sequencing limitations (64,65). Studies of plasmid transmission in natural populations are rare, though others have compared gene content and plasmid diversity among non-coexisting plasmids within a single species (60,66). Here we establish a novel framework for studying plasmid transmission in nature and find quite different patterns of inheritance among the multiple plasmids that coexist within *Rhizobium* host cells.

Some plasmids appear to move more vertically alongside the chromosome, while others move horizontally. Given that these plasmids co-occur in the same cells, plasmid-specific mechanisms must explain this variation, at least in part. Our findings support evolutionary conceptions of plasmids as having their own agency and fitness interests (61,67,68).

Rhizobia also provide a rare opportunity to study how plasmids and HGT facilitate quantitative trait evolution in natural bacterial populations. Rhizobial symbiosis genes are known to be both horizontally-transmitted and selected in nature (34,35). In our *Rhizobium* population, we find frequent HGT of the pSym relative to the other plasmids paired with differentiation of the pSym between N environments, suggesting environmentally-dependent selection on this plasmid. The symbiosis genes in *Rhizobium* can move between plasmids (32,35,38,39), and as we show here, those plasmids can vary in rates of HGT – potentially suggesting that the traits governed by these mobile loci will evolve more or less rapidly depending on their genomic location. Rapid evolution of symbiotic traits via gene-specific sweeps, whole plasmid sweeps, and even plasmid loss (69) might be advantageous in context dependent mutualisms where the costs and benefits of symbiosis shift with the biotic and abiotic context in which partners interact (70,71).

By integrating genomics, plasmid biology, phylogenetics, and plant-microbe interactions in wild bacteria, our study provides a framework for quantifying the relative rates of vertical and horizontal transmission among the ECEs that coexist within a single species and elucidates the role of plasmid HGT in an ecologically-important symbiosis that plays a critical role in global nitrogen cycling.

## MATERIALS AND METHODS

### Strain isolation and growth

Here we generate novel genomes for a population of 62 *Rhizobium* strains originally isolated from old field successional plots at the KBS LTER. Full methods detailing the long-term N fertilization experiment, strain isolations, and phenotypic experiments to characterize partner quality symbiosis with three clover host species are described elsewhere (41). Briefly, rhizobium strains were isolated from both N-fertilized and control plots. Fertilized plots had been supplemented with 12.3 g N m^−2^ per year granular ammonium nitrate for 22 years prior to sampling, whereas control plots remained unfertilized.

### DNA and sequencing

We grew the strain isolates in solid tryptone yeast (TY) media (5 g L^-1^ tryptone, 3 g L^-1^ yeast extract, 6 mM CaCl2, and 16 g L^-1^ agar) plates at 30 C for 2 days. After growth on solid media, single colonies were selected to inoculate 5 mL liquid TY media for 1 day at 30 C in a roller drum. The PacBio Nanobind CBB kit (Pacbio, San Diego CA) was used to extract high molecular weight (50–300+ kbp) DNA from 1 mL of bacterial culture for all strain isolates. The DNA was sent for PacBio hifi long-read sequencing (72) at the W. M. Keck center at the University of Illinois, where gDNAs were sheared with a Megaruptor 3 to an average fragment length of 10kb then converted to barcoded libraries with the SMRTBell Express Template Prep kit 3.0 and pooled in equimolar concentration. The pooled libraries were sequenced on 2 SMRTcell 8M on a PacBio Sequel IIe using the CCS sequencing mode and a 30hs movie time. Circular consensus sequence (CCS) analysis was done using SMRTLink V11.0 using the following parameters: ccs --min-passes 3 --min-rq 0.99 lima --hifi-preset SYMMETRIC --split-bam-named --peek-guess.

### Genome assembly and annotation

All genomes were assembled using recommended workflows in Trycycler (73). Briefly, raw reads were filtered for quality using Filtlong, assessing both length and quality of the reads. The raw reads where then divided into a 12 maximally-independent subsets using the subsample function in Tryclycler. Next Trycycler uses three assembly methods (Flye (74), Hifiasm (75), and Raven (76)) to generate independent whole-genome assemblies for four read subsets each. Finally we used Trycycler to generates a consensus genome based on these 12 assemblies, followed by manual curation to ensure consistent genome structure across assemblies. Genomes were then annotated using NCBI’s Prokaryotic Genome Annotation Pipeline (PGAP).

### Classification of plasmids

We generated individual fasta files for every replicon in every strain then used sourmash (77) to generate and compare k-mer signatures and calculate pairwise Jaccard Index (JI) values using the parameters: sketch dna -p scaled=10000, k=31, compare -p 8. Signature tables were then imported into Cytoscape, and component graphs were created using a minimum JI value of 0.1 to delineate clusters; this cutoff was chosen based on similar analyses of *Rhizobium*, *Agrobacterium*, *Bradyrhizobium*, as well as a global plasmid analysis (7,37,78).

We classified the resulting clusters as plasmid “types”, then layered this plasmid presence onto phylogenetic trees using “ggtree” and “ggplot” libraries in R (version 4.3.2 “eye holes”). We used the package popgenome to calculate pi and Average Nucleotide Identity (ANI) for aligned core regions of each plasmid type.

### Phylogenetic trees

We used a custom SPINE-Nucmer pipeline (https://github.com/Alan-Collins/Spine-Nucmer-SNPs) to generate core genome alignments for all 62 genomes (core genes were all chromosomal), all chromosomes (resulting in the same core), and subsequently for each plasmid type. For each, core components were concatenated, aligned using MAFFT, and used to generate phylogenetic trees in IQTree2 with parameters: -bb 10000 -st DNA.

### Presence-absence data and Principal Coordinate Analysis

We used PIRATE (79) on PGAP-annotated genomes to define orthologous gene presence-absence using PIRATE (79)., followed by Principal Coordinate Analysis (PCoA) of this output using the “dplyr”, “vegan” packages in Rstudio to calculate Jaccard distances between all samples based on shared gene content. Pairwise distances were then transformed using the “cmdscale” function from the “stats” package and sketched using the “ggplot” package. Gene presence-absence plots were made using “pheatmap” package.

### Gene-tree simulations

To assess patterns of plasmid inheritance, we created custom scripts to generate distributions of gene tree distances among genes randomly subsampled from within (or across) replicons in our population – with larger gene tree distances indicating more gene tree heterogeneity and thus increased horizontal (versus vertical) transmission. First we generated a list of the core genes in each plasmid using PIRATE. For each replicon, we used custom R scripts to randomly subsample two sets of 100 genes (10 from the small core of the type IV/pSym) from each strain, align them, create two phylogenetic trees, and calculate Generalized Robinson-Foulds (GRF) distance between trees, then repeated this process over 1000 random resamplings. We plotted these GRF distributions using geom_density function in the ggplot2 package. Next we repeated this process comparing samples from each plasmid to the chromosome. Because the chromosome is necessarily vertically transmitted each generation, the distribution of chromosome-to-chromosome comparisons serves as a null expectation for the GRF distribution of a vertically inherited element in the presence of horizontal gene transfer, recombination, and gene tree uncertainty.

## Availability of data and materials

Phenotypic data, and scripts used for generating analyses will be available upon acceptance. Genomes will be uploaded to NCBI genome database upon acceptance.

## ACKNOWLEDGMENTS

D.V.G., K.D.H., C.K.V., and R.J.W. conceived the project. D.V.G. extracted and submitted DNA for sequencing. D.V.G. and C.P.S. generated the genomes and performed bioinformatic analyses. D.V.G. and K.D.H. drafted the article, and all authors participated in critical revisions and approved the final version for submission. K.D.H., C.K.V., R.J.W., J.A.L. acquired funding for the project.

We would like to acknowledge Dr. Alvaro Hernandez and Chris Wright from the Roy J. Carver Biotechnology Center at the University of Illinois for library preparation and sequencing. We thank Dr. Jaya Chandrashekhar, Dr. Susan Thomas, and the rest of GEMS for their feedback, help, and coordination of lab protocols. We thank Dr. Ilan Shomorony, Dr. Pamela Martinez, and Ivan Sosa-Marquez for their invaluable help, motivation, and insightful advice during the manuscript writing process. The results and interpretation of this manuscript accomplished by J.G.M. were in their personal capacity and are the author’s own views and do not reflect the view of their employer.

## Competing interests

The authors declare that they have no competing interests

## Funding

This research is a contribution of the GEMS Biology Integration Institute, funded by the National Science Foundation DBI Biology Integration Institutes Program, Award # 2022049, as well as NSF Award #1257938.

---

**Figure S1:**
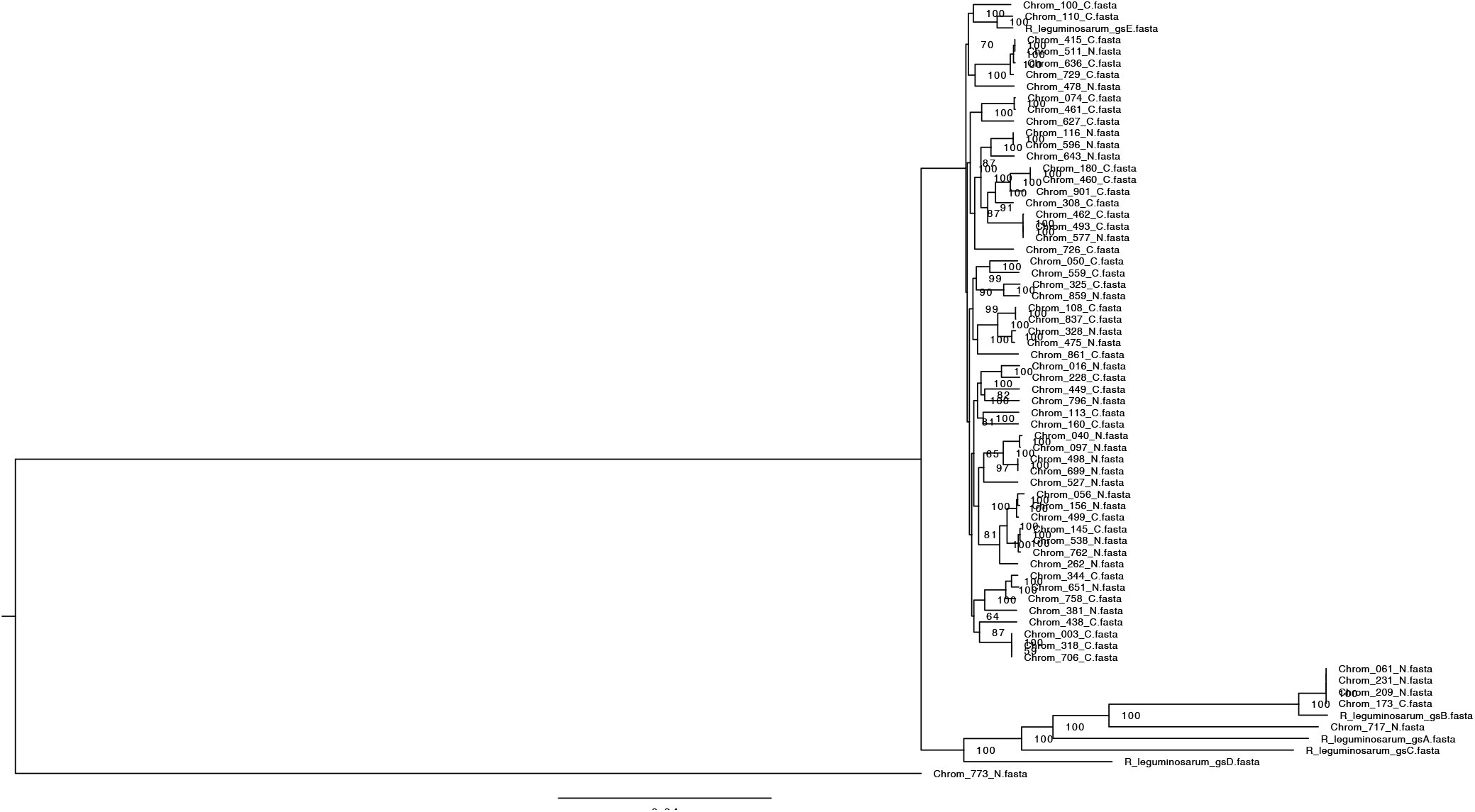
Chromosomal phylogeny of clover-associated *Rhizobium* strains from the KBS LTER population alongside one representative each from five genospecies (A-E) of the *Rhizobium* species complex, as described in Cavassim et al. 2020. Tree was manually rooted on strain 717_N. Any nodes with < 85% bootstrap support were collapsed.

**Figure S2:**
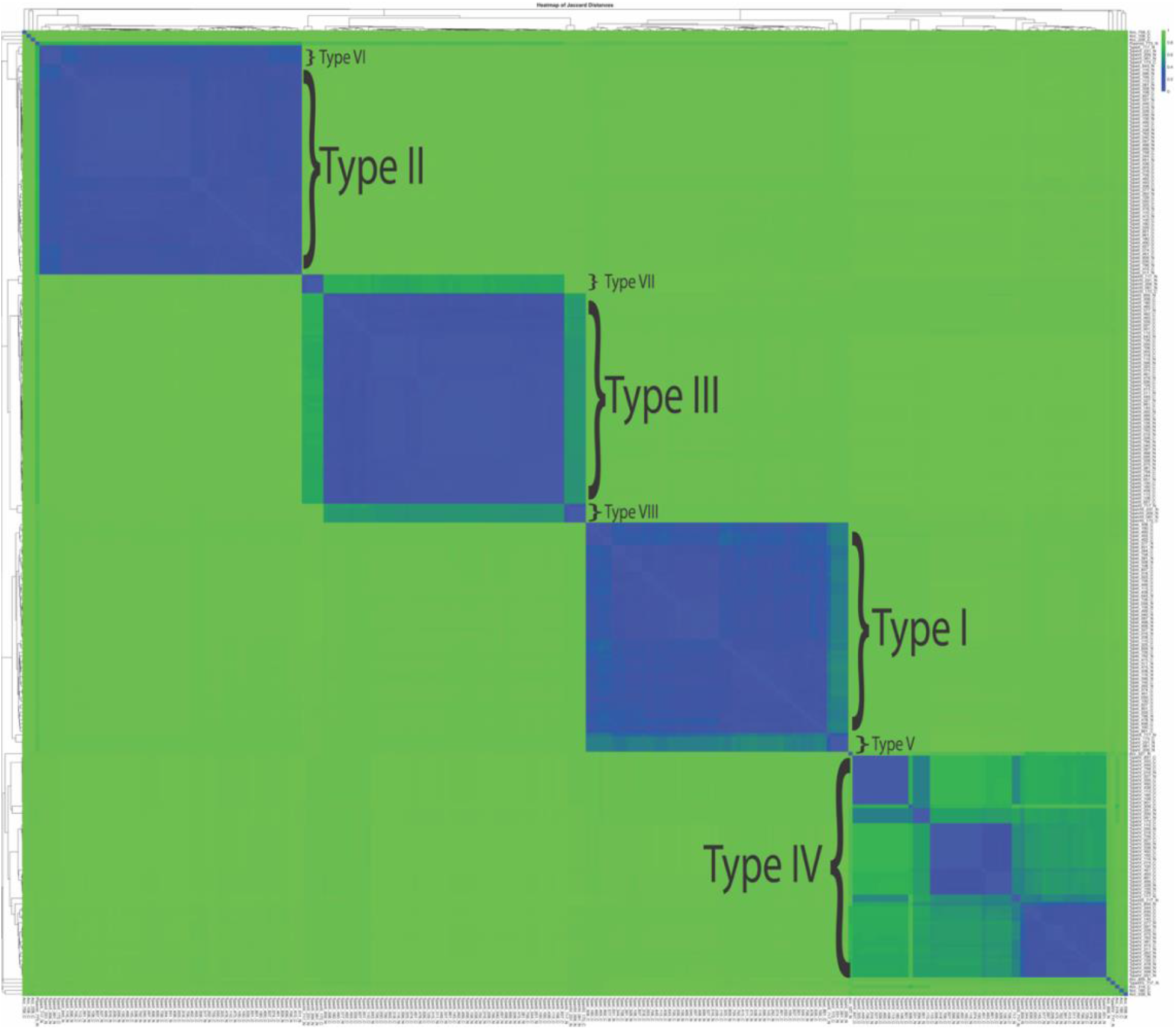
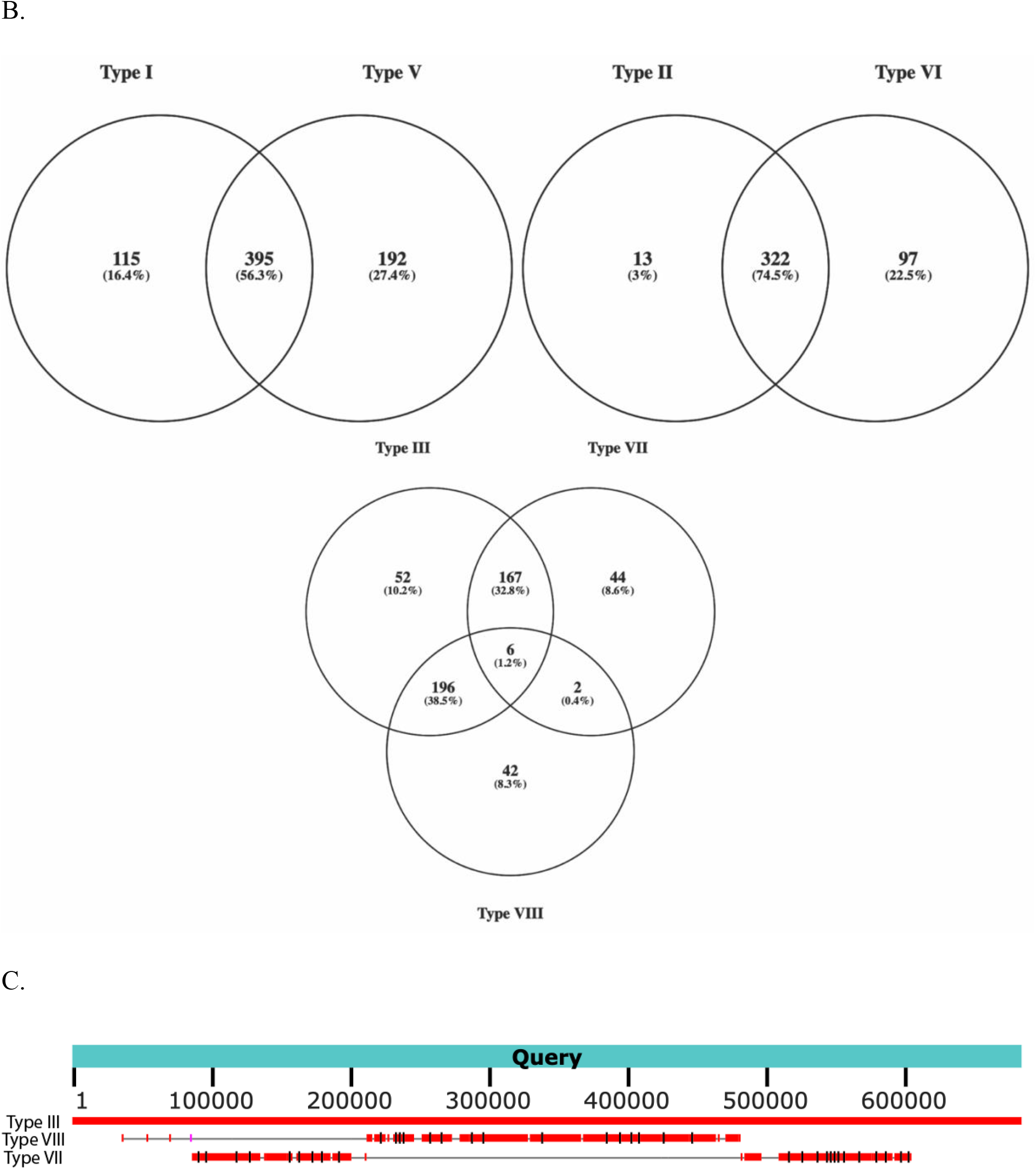
(A) Heatmap of pairwise Jaccard distances based on gene presence-absence between all plasmids from our study, highlighting similarity in gene content between plasmid types from *gsE* and *gsB*: I with V, II with VI, and III with VII/VIII. (B) Venn diagram of genes shared across plasmid types III, VII, and VIII. (C) local alignment of a representative of type III (strain 110_C) with representatives of types VII and VIII (both from 231_N).

**Figure S3:**
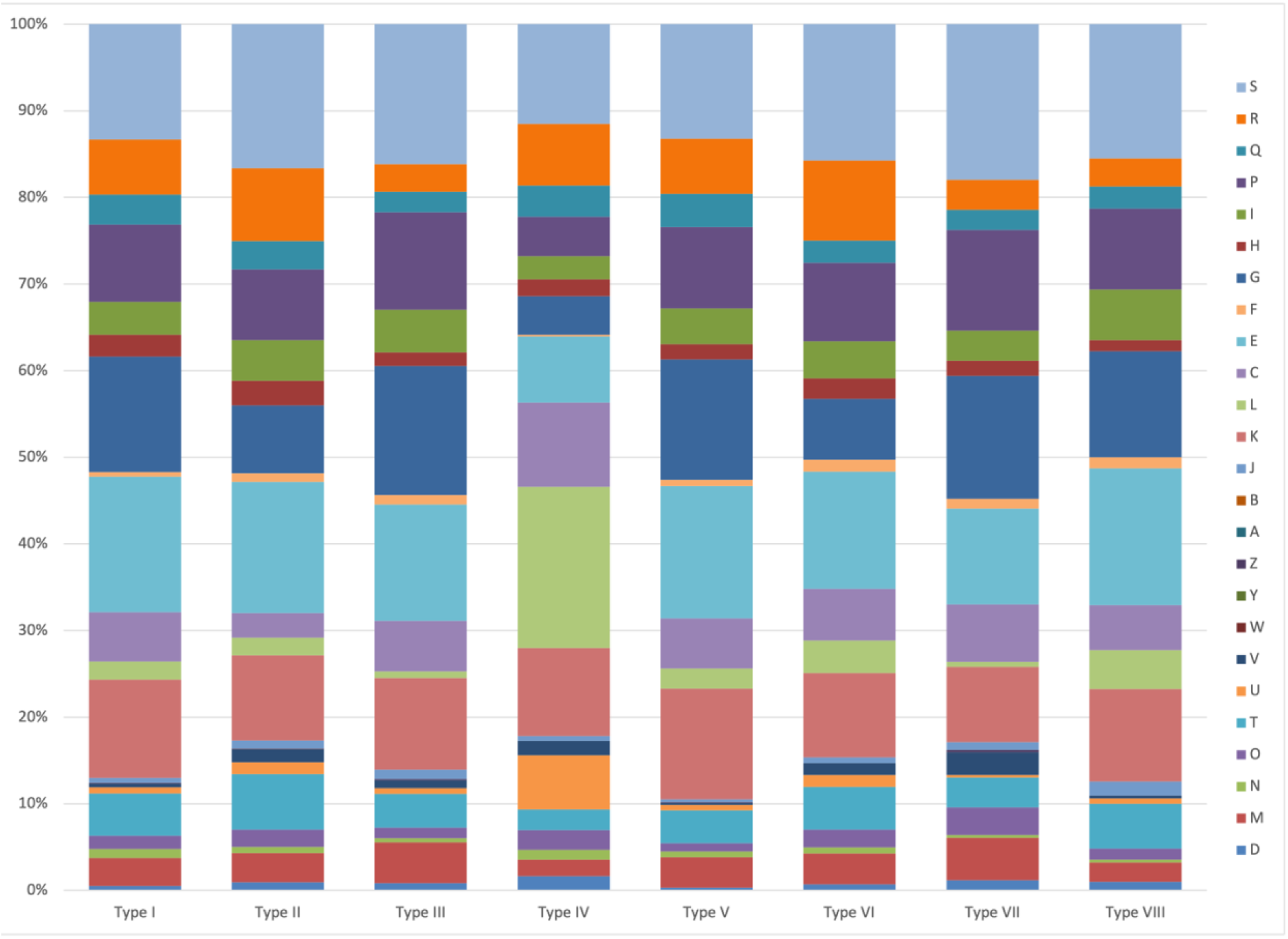
Distribution of COG functions across the plasmids of clover-associated *Rhizobium* from a natural population: (S) Function unknown, (R) General function prediction only, (Q) Secondary metabolites biosynthesis, transport and catabolism, (P) Inorganic ion transport and metabolism, (I) Lipid transport and metabolism, (H) Coenzyme transport and metabolism, (G) Carbohydrate transport and metabolism, (F) Nucleotide transport and metabolism, (E) Amino acid transport and metabolism, (C) Energy production and conversion, (L) Replication, recombination and repair, (K) Transcription, (J) Translation, ribosomal structure and biogenesis, (B) Chromatin structure and dynamics, (A) RNA processing and modification, (Z) Cytoskeleton, (Y) Nuclear structure, (W) Extracellular structures, (U) Intracellular trafficking, secretion, and vesicular transport, (T) Signal transduction mechanisms, (O) Posttranslational modification, protein turnover, chaperones, (N) Cell motility.

**Figure S4:**
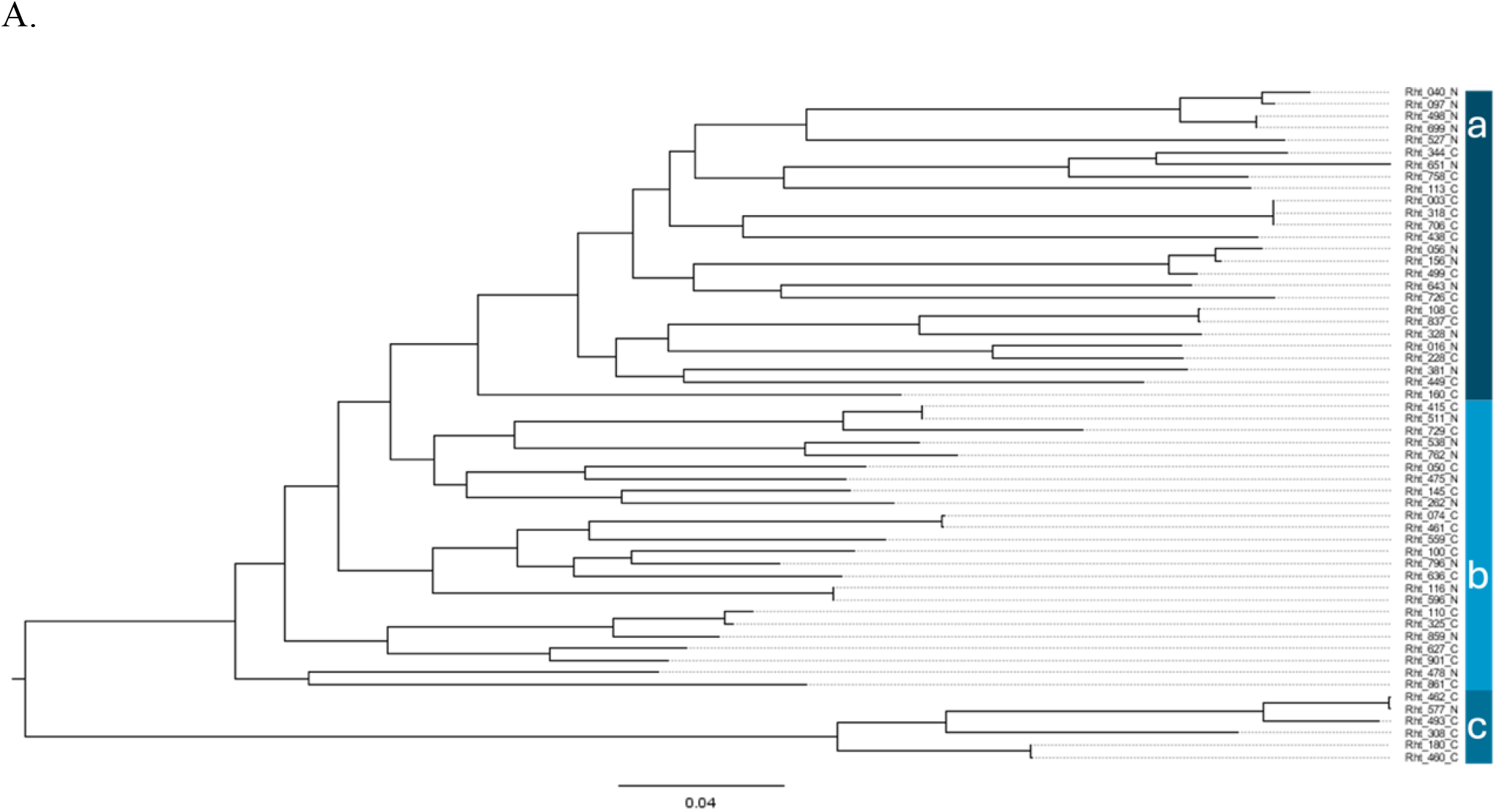
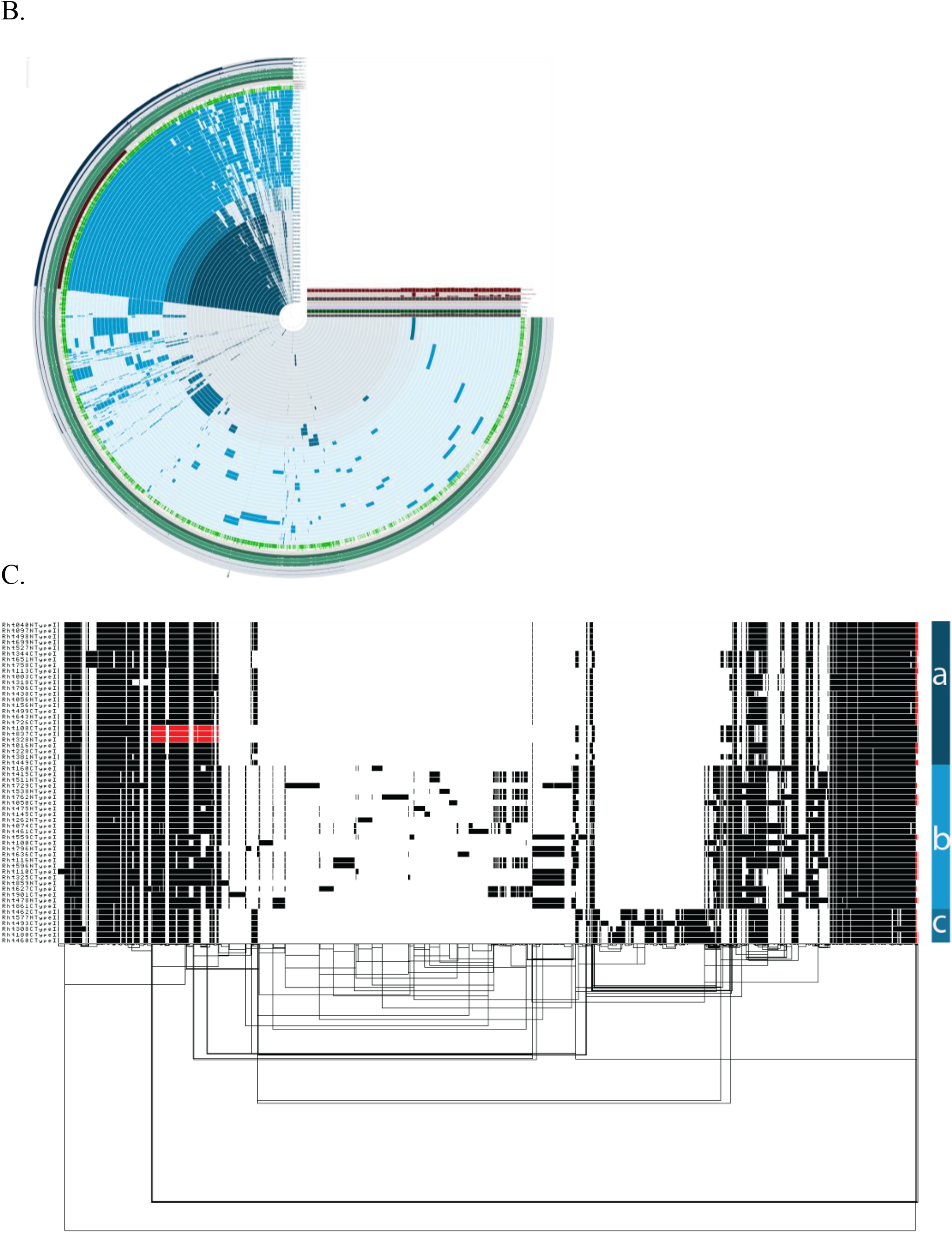
(A) Phylogenetic tree of the concatenated core genes of the type I plasmid, showing the relationships among the three subclades clades as in Figure 1. Any nodes with < 85% bootstrap support were collapsed. (B) Gene presence-absence anv’io plot of Type I plasmids, colored by clade and showing that larger subclades I-b and I-c possess distinct large insertions. (C) Pangenome graph view of the Type I plasmid population, showing distinct insertions in larger subclades I-b and I-c occur in the same genomic location. Aligned sequences are in black, and inverted regions are in red.

**Figure S5:**
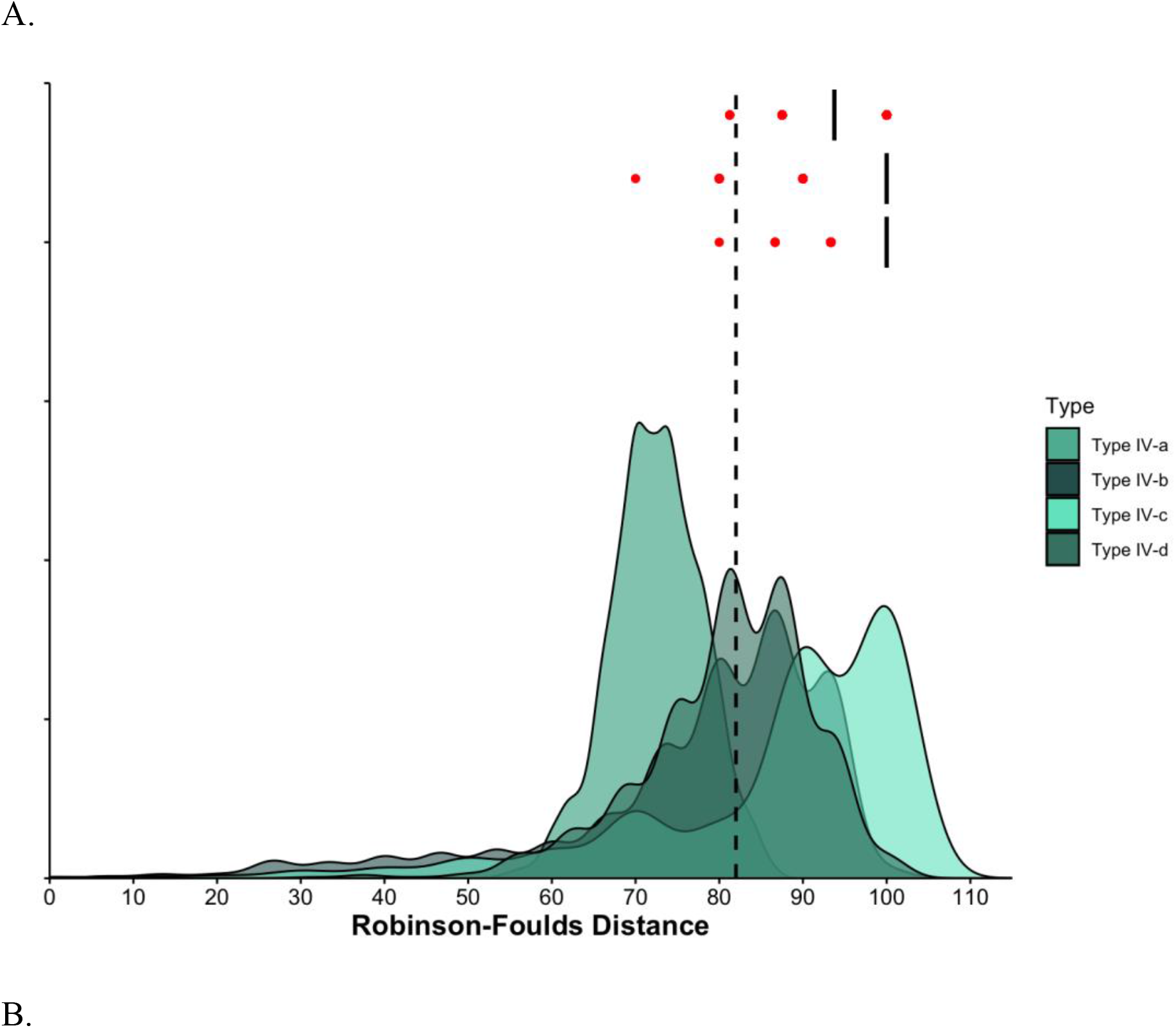
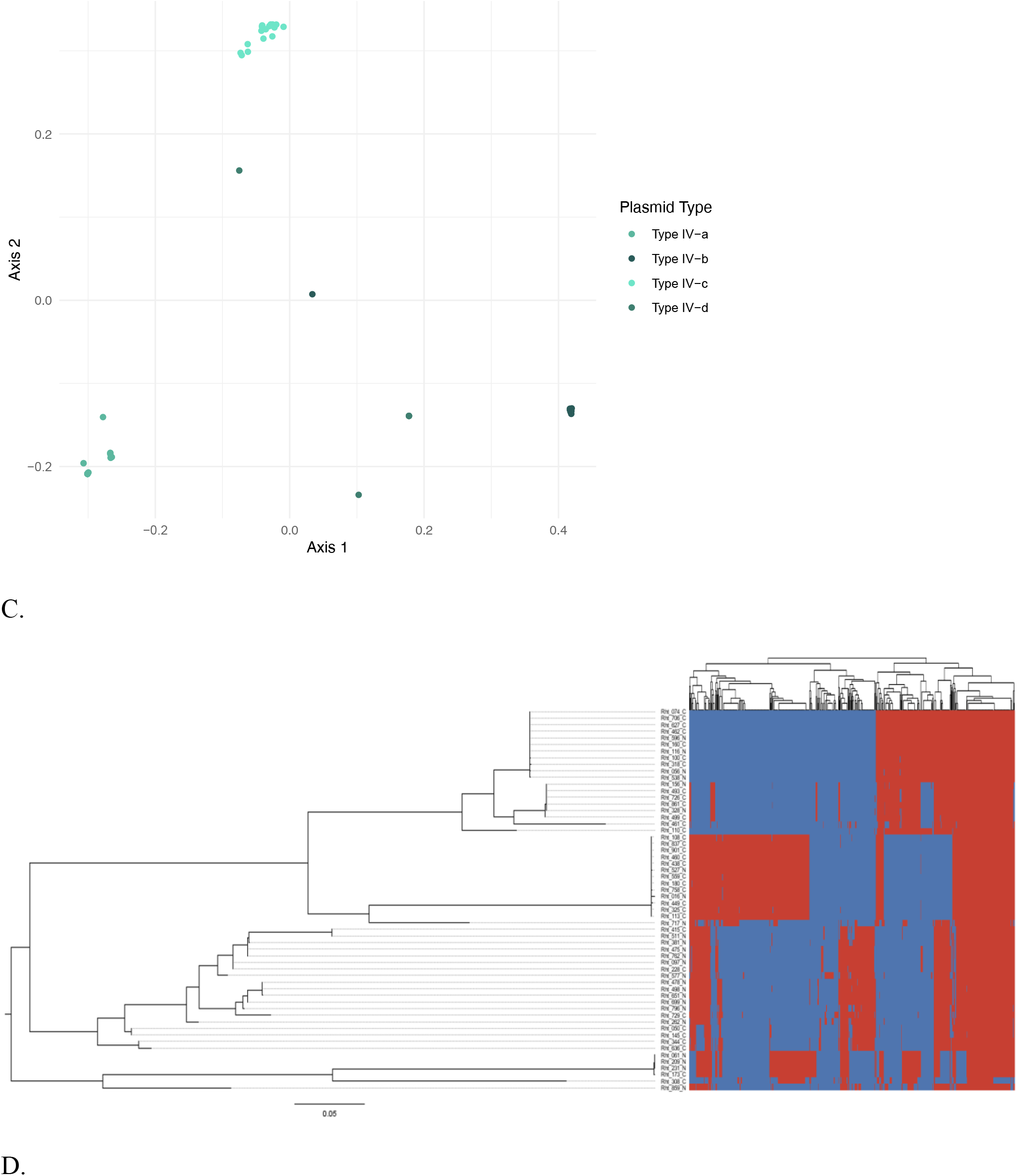
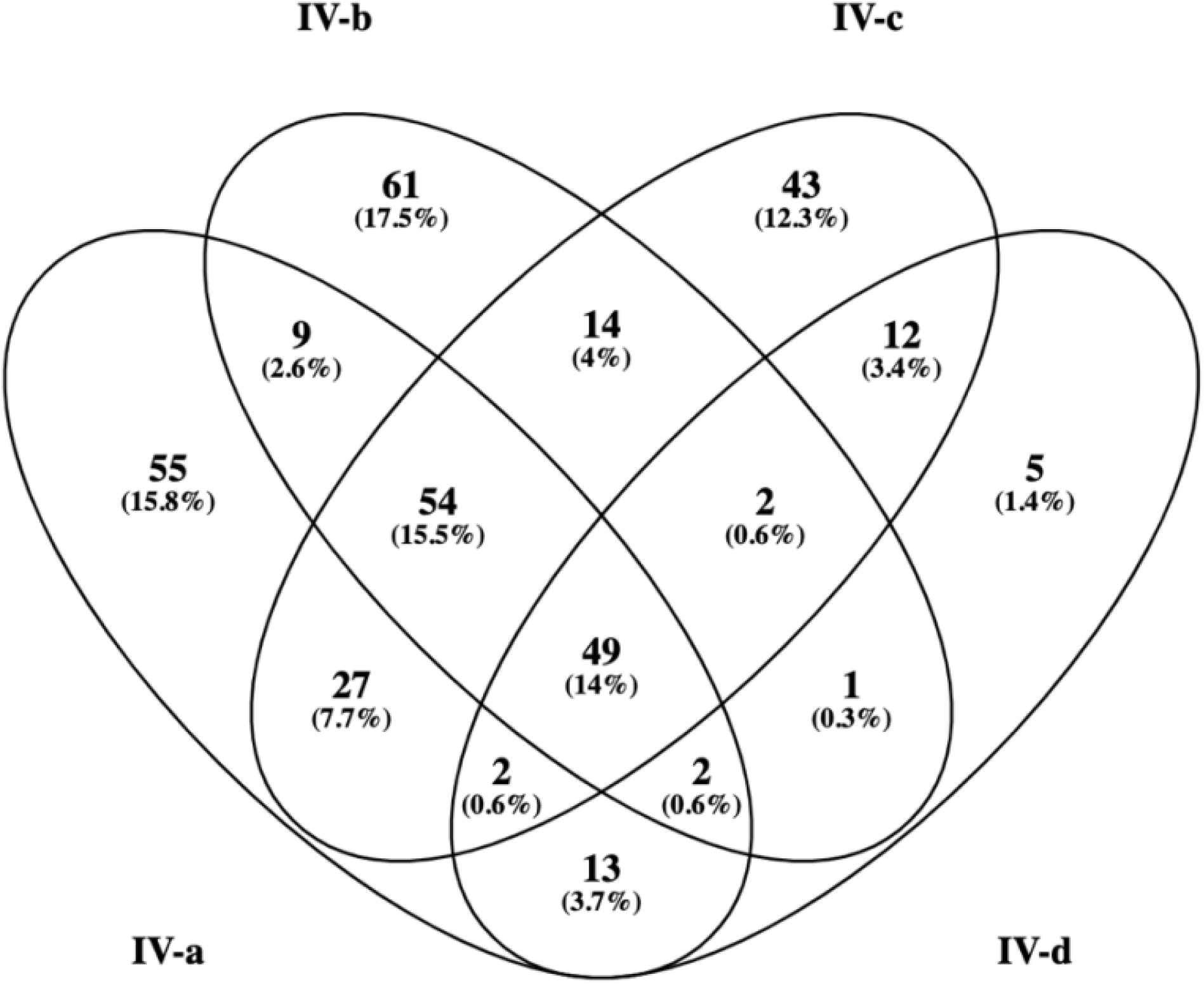
(A) GRF distances within Type IV subclades (smoothed distributions) and between subclades and the chromosome (box and whisker plots). (B) Principal Coordinate Analysis (PCoA) of orthologous gene clusters in the type IV plasmids in *Rhizobium.* (C) Tree of type IV pSym showing orthologous genes that are present (red) or absent (blue) across strains. (D) Venn diagram showing the number of core genes in different combinations of type IV subclades.

**Table S1:** Size and strain of origin of accessory plasmids in the pangenome.

| Accessory plasmid's strain | Size (bp) |
| --- | --- |
| Rht_108_C | 72609 |
| Rht_180_C | 122968 |
| Rht_228_C | 73852 |
| Rht_318_C | 188843 |
| Rht_527_N | 277757 |
| Rht_538_N | 245492 |
| Rht_758_C | 22509 |
| Rht_859_N | 208987 |

